# Cell surface localization and intercellular transfer of the hantavirus nucleocapsid protein via a viral assembly-independent mechanism

**DOI:** 10.64898/2025.12.05.692699

**Authors:** Amelina Albornoz, Natalia Salazar-Quiroz, Eduardo A. Bignon, Nicolás A. Muena, Diego Gonzalez, Jarell Espinoza, María José Fuenzalida, Marcelo López-Lastra, Rainer G. Ulrich, Nicole D. Tischler

**Affiliations:** Laboratorio de Virología Molecular, Centro Científico Tecnológico de Excelencia Ciencia & Vida, Fundación Ciencia & Vida, Santiago, Chile; Escuela de Medicina, Facultad de Medicina, Universidad San Sebastián, Santiago, Chile; Escuela de Bioquímica, Facultad de Ciencias, Universidad San Sebastián, Santiago, Chile; Laboratorio de Virología Molecular, Instituto Milenio de Inmunología e Inmunoterapia, Departamento de Enfermedades Infecciosas e Inmunología Pediátrica, Escuela de Medicina, Facultad de Medicina, Pontificia Universidad Católica de Chile, Santiago, Chile; Institute of Novel and Emerging Infectious Diseases, Friedrich-Loeffler-Institut, Federal Research Institute for Animal Health, Greifswald-Insel Riems, Germany

**Author notes:** Correspondence to Nicole D. Tischler,;. Yale School of Medicine, Department of Internal Medicine, Section of Infectious Diseases, New Haven, USA. Institut Pasteur, Unité de Virologie Structurale, Departement de Virologie, Paris, France (Eduardo A. Bignon).

## Abstract

Hantaviruses are segmented negative-sense RNA viruses that package their genome into helical ribonucleocapsids, by oligomerization of the nucleocapsid (N) protein along the viral RNA. Upon transmission to humans, hantaviruses can cause severe disease characterized by increased vascular permeability in the microvasculature, a key driver of pathogenesis. In patients and rodent reservoirs, robust and persistent antibody responses against the nucleocapsid (N) protein are consistently observed. Here we investigated whether the hantavirus N protein, described as a strictly intracellular protein, is released into the extracellular environment during viral infection.

By using diverse experimental approaches, we demonstrate that in cells infected with Andes virus (ANDV) or the non-pathogenic Tula virus (TULV), the N protein localizes on the plasma membrane. Transfection experiments further revealed that the surface localization of N proteins from *Murine*, *Arvicoline*, *Neotomine* and *Sigmodontine* rodent-reservoir-associated hantaviruses did not require co-expression of other viral proteins indicating that its trafficking to the cell surface is mediated by a mechanism distinct from canonical virion assembly pathways. Moreover, we show that the N protein is secreted from cells into the extracellular milieu and is further transferred to neighboring cells. Interestingly, despite being present on the cell surface and secreted into the extracellular media, the TULV N protein transfer of the N protein to neighboring cells was significantly lower compared to that of ANDV N protein. The extracellular presence of hantavirus N proteins, and likely also from other clinically-relevant bunyaviruses, unveils new possible functions of the protein during the viral infection cycle, which is likely related to immune responses, immune modulation and viral pathogenesis.

## INTRODUCTION

Rodent host-associated hantaviruses from the *Orthohantavirus* genus can cause severe hantavirus disease in humans; however, no effective antiviral countermeasures are currently available [1, 2]. Puumala virus (PUUV) and Hantaan virus (HTNV) in Europe and Asia can cause hemorrhagic fever with renal syndrome (HFRS) while Andes virus (ANDV) and Sin Nombre virus (SNV) in South and North America can lead to severe hantavirus cardiopulmonary syndrome (HCPS). Furthermore, for ANDV human-to-human transmission cascades have been reported, generating pandemic awareness [3–5]. To date, no effective treatment against hantaviruses nor vaccine against HCPS-causing hantaviruses are available. Some hantaviruses have only been associated with human disease in sporadic cases such as Tula virus (TULV) in Eurasia [6] and have no association in the case of Prospect Hill virus in North America [7].

Hantaviruses which belong to the *Bunyaviricetes* class, are enveloped viruses with a tri-segmented negative-sense (-) single-stranded (ss) RNA genome encoding four structural proteins. The surface glycoproteins Gn and Gc form hetero-octameric spikes and mediate viral cell entry and egress [8–10]. The genomic RNA is encapsidated by the N protein, while the RNA-dependent RNA polymerase (RdRp) binds the 5′ and 3′ RNA termini to form the viral ribonucleocapsids [11–13]. N protein drives genome encapsidation via multimerization along viral RNA (vRNA) through a positively charged groove [12] and also participates in vRNA replication, RNA transcription via cap-snatching, and mRNA translation [14–18]. During replication, N is the most abundant viral protein and is detectable 2-4 hours post infection (hpi). N appears in small cytoplasmic puncta and later forms filamentous structures [19]. An early report evidenced that overexpression of N alone leads to the production of ribonucleoprotein-like structures (RLPs) within cells [20]. Within cells, the N protein also associates with intracellular membranes near the Golgi apparatus, likely corresponding to replication factories containing vRNA, mRNA, and N protein [20–23]. It also co-localizes with stress granules and P-bodies where cap-snatching may occur [15, 23–25]. However, not all punctate N protein staining co-localizes with RNA or stress granules [23, 25], re-enforcing its multifunctionality [17]. N protein also inhibits apoptosis; for TULV N protein it has been shown to sequester activated caspase-3 [26] while in the case of ANDV, Dobrava-Belgrade virus and PUUV, the N protein blocks granzyme B activity, which counteracts lymphocyte-mediated killing [27, 28]. Additionally, in microvascular endothelial cells expressing ANDV N protein under hypoxic conditions, it increases cellular permeability [29]. Moreover, the N protein of ANDV, unlike N from other hantaviruses, inhibits the cellular antiviral innate immune response by preventing TBK-1 autophosphorylation and targeting IFN-β induction [30].

In patients, strong and long-lasting antibody responses against the N protein are observed, detectable from the acute phase and persisting for years [31–33]. They mostly target the first 100-120 amino acids of the N-terminal region are highly antigenic [34, 35] although other regions also present immunogenicity in human and rodents [36]. The immunization with the N protein elicits specific antibodies 10 days post-infection and can induce a protective immunity in natural reservoir animals [35, 37–45]. Overexpression of Gn, Gc, and N protein induces virus-like particle (VLP) release, though the glycoprotein precursor (GPC) alone can also form and secrete VLPs in heterologous expression systems [20, 46, 47]. The strong, early and immunodominant humoral immune response against the N protein in human and rodent hosts infected with different hantaviruses [32, 33], raises the question of whether during viral infection, the N protein may be exposed to the immune system in the extracellular space. Although cell surface accumulation of the N protein has been reported for several segmented and non-segmented (-)ssRNA viruses, including influenza A virus, vesicular stomatitis virus [48, 49], severe acute respiratory syndrome (SARS)-coronavirus (CoV)-2 [50], measles virus [51] and respiratory syncytial virus (RSV) [52], to date there is no evidence that supports the extracellular exposure of the N protein of hantaviruses or other bunyaviruses.

Through multiple experimental approaches, this study reports on the detection of the N protein of several hantaviruses on the surface of various cell lines, both in the presence and absence of other viral components. The data further show that the N protein from several hantaviruses is secreted into the extracellular medium. Notably, the N protein of ANDV was efficiently transferred to neighboring cells, while significantly less transfer occurred with N protein from TULV. Collectively, these findings reveal previously unrecognized trafficking, secretion, and intercellular transfer properties of the hantavirus N protein, thereby broadening our understanding of fundamental viral infection properties and its interaction with the host ultimately providing a framework for the development of novel antiviral strategies.

## MATERIALS AND METHODS

### Cells and viruses

Human endothelial EA.hy926 (American Tissue Culture Collection (ATCC) number CRL-2922) and epithelial Vero E6 (ATCC number CRL-1587) cells were grown in modified Eagle medium (MEM) with 10% fetal bovine serum (FBS, GIBCO), 1x essential amino acids (Thermo Fisher Scientific) and 1 mM sodium pyruvate (Thermo Fisher Scientific). Human embryo kidney HEK293FT cells (Thermo Fisher Scientific) were propagated in Dulbeccós MEM (DMEM) supplemented with 10% FBS (GIBCO). All work involving the infectious ANDV isolate CHI-7913 (*Orthohantavirus andesense*) [53] (kindly provided by TM Héctor Galeno, Instituto de Salud Pública, Chile) was performed under strict biosafety level three (BSL3) conditions (Centro de Investigaciones Médicas, Pontificia Universidad Católica de Chile, Chile). The work with TULV isolate Moravia/5302Ma/94 strain (*Orthohantavirus tulaense*) [54] (kindly provided by Drs. Detlev H. Krüger and Boris Klempa, Charité Universitätsmedizin Berlin, Germany) was performed under BSL2 conditions. Virus titers were quantified by flow cytometry as described below, only after viral inactivation achieved by incubating the cells during 30 min with 2% paraformaldehyde (PFA).

### Antibodies

The monoclonal antibodies (MAb) used in this study are summarized in Table.

**Table 1.**
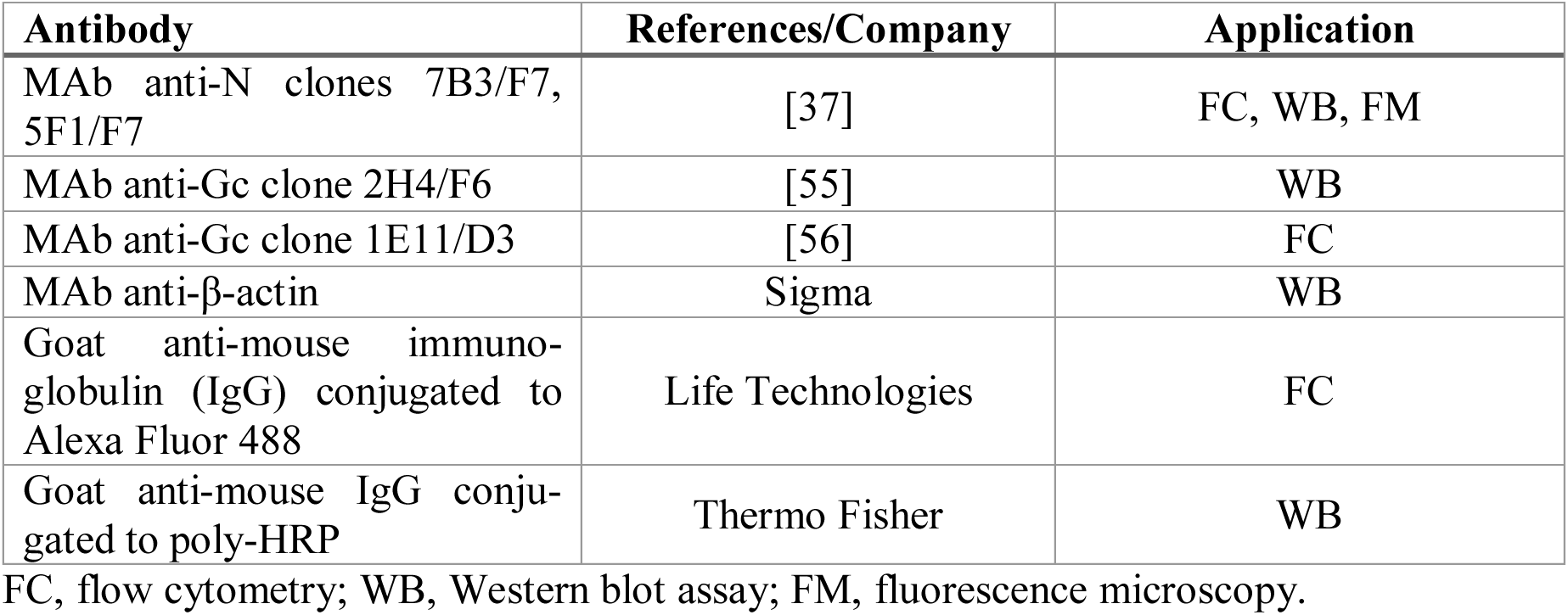
Antibodies used in this study and their applications.

| Antibody | References/Company | Application |
| --- | --- | --- |
| MAb anti-N clones 7B3/F7, 5F1/F7 | [37] | FC, WB, FM |
| MAb anti-Gc clone 2H4/F6 | [55] | WB |
| MAb anti-Gc clone 1E11/D3 | [56] | FC |
| MAb anti-β-actin | Sigma | WB |
| Goat anti-mouse immuno-globulin (IgG) conjugated to Alexa Fluor 488 | Life Technologies | FC |
| Goat anti-mouse IgG conjugated to poly-HRP | Thermo Fisher | WB |

FC, flow cytometry; WB, Western blot assay; FM, fluorescence microscopy.

### Plasmids

As an empty plasmid we used pI.18 (kindly provided by Dr. Jim Robertson, from the National Institute for Biological Standards and Control, Hertfordshire, UK). For ANDV Gn/Gc expression we used the plasmid pI.18/GPC that encodes for the Gn/Gc proteins as full-length GPC of ANDV strain CHI-7913 [40]. For the expression of the N protein from different hantaviruses, we used pCMV-Bios/N-ANDV encoding the entire N protein of ANDV strain CHI-7913 [33], p/HA-SNV-N encoding full-length N protein from SNV strain SN77734 [57] and prepared the vectors pcDNA3/N-Tula95s encoding full-length N protein of TULV strain Moravia (GenBank accession number Z69991 [54]) and pcDNA3-VR1 encoding full-length N protein of PUUV strain Vranica/Hällnäs (GenBank accession number U14137) [58]. The entire TULV and PUUV N protein coding sequences were generated by RT-PCR using infected Vero E6 cells, cloning of the RT-PCR products into pCR-TOPO vectors, re-sequenced and finally subcloned into pcDNA3 (Invitrogen) plasmid using methods indicated elsewhere [59].

### Confocal immunofluorescence microscopy of infected cells

For confocal microscopy analysis, Vero E6 cells were seeded on coverslips in 24-well plate with MEM medium and 10% FBS overnight. Cells were infected with ANDV (multiplicity of infection, MOI 1) for 1 h at 37°C. After 24 h, cells were incubated with CellMask^TM^ Orange plasma membrane stain (Thermo Fisher Scientific) for 10 min at 37°C to prevent internalization, and next fixed in 4% PFA for 15 min at room temperature to inactivate the virus. For immunofluorescence, permeabilized cells were incubated in 0.1% Triton X-100, followed by incubation with the primary antibody. In contrast, non–permeabilized cells were directly incubated with the primary antibody. MAb anti-N protein clone 7B3/F7 was used at a dilution of 1:2,000 for 1h at 37°C. Secondary goat anti-mouse IgG conjugated to Alexa Fluor 488 was used at a dilution of 1:500 (Invitrogen A11001). The samples were incubated with 4′,6-Diamidin-2-phenylindol (DAPI; 3 nM, Thermo Fisher Scientific) for 5 min at room temperature. Z-stack confocal images were acquired with a Zeiss Airyscan confocal microscope using a 63X objective (Unidad de Microscopía Avanzada, Pontifícia Universidad Católica de Chile). Image analyses and orthogonal views were generated with Fiji software (ImageJ, version 2.14.0/a.54f) to visualize spatial distribution of the N protein in infected cells.

### Flow cytometry analysis of infected or transfected cells

For viral infection, Vero E6 or EA.hy926 cells grown in 24-well plates were incubated for 1 h at 37°C with ANDV or TULV (MOI 1) and the infection was stopped by detaching the cells with trypsin and fixing with PFA after the indicated time points. For plasmid DNA transfection, HEK293FT cells were seeded in 100 mm dishes and 24 h later transfected with 8 μg of plasmid DNA using the calcium phosphate protocol [60]. Vero E6 and EA.hy926 cells were seeded in 24-well plates and transfected the next day with 0.5 μg of plasmid DNA per well using Lipofectamine 2000 (Invitrogen). Viral infection or plasmid DNA transfection was quantified at the indicated time points by detecting cells expressing the viral N protein by use of flow cytometry as previously described [61]. Briefly, before fixation, infected or transfected cells were incubated for 15 min with the Live/Dead Fixable Violet Dead Cell Stain Kit (Live/Dead staining; Thermo Fisher Scientific) as described by the manufacturer to analyze only live cells. Next, infected or transfected cells were fixed for 30 min with 2% PFA and subsequently used either without permeabilization to detect cell surface proteins or after permeabilization to detect the total amount of proteins of cells. For immunostaining, the cells were incubated overnight at 4°C with anti-N MAb (5F1/F7: 1:1,000, 7B3/F7 same dilution as described above) or anti-Gc Mab (clone 1E11/D3). After washing, they were labeled with goat anti-mouse IgG conjugated to Alexa Fluor 488 (1:1,000) (Thermo Fisher Scientific). Flow cytometry was performed in a cytometer (FACS Canto II, Becton Dickinson). The gate for ANDV N-positive, live cells was established using the Live/Dead staining and a negative control consisting either of non-infected cells or cells transfected with the pI.18 empty control plasmid (Mock), which were incubated with the same primary and secondary antibodies.

### Biotinylation

For GPC or N expression, HEK293FT cells were grown in 100 mm plates and calcium phosphate-transfected with the corresponding plasmid DNAs. Forty-eight h later, cell surface proteins were labeled with biotin in order to separate the biotinylated (surface proteins) from non-biotinylated (intracellular proteins) fractions using a cell surface protein isolation kit (Pierce). In addition, the protein content in the supernatant was concentrated by precipitation using ammonium sulfate at a final concentration of 35% (w/v) [62]. For protein detection by western blot assay, primary MAb anti-N clone 7B3/F7 was used at a dilution of 1:2,000, MAb anti-Gc clone 2H4/F6 or anti-β-actin MAb (Sigma) were used at 1:2,500 dilution and subsequently detected with an anti-mouse IgG horse radish peroxidase (HRP) conjugate (Thermo Fisher Scientific) at a dilution of 1:5,000 and a chemiluminescent substrate (Super Signal^TM^ West Pico PLUS Chemiluminescent Substrate, Thermo Fisher Scientific).

### N protein cell transfer assay

EA.hy926 cells were seeded overnight on coverslips in 24-well plates in MEM medium containing 10% FBS. The next day, cells were transfected with plasmid DNAs pCMV-Bios/N-ANDV or with pLVX-mCherry-c1 using Lipofectamine 2000 (Thermo Fisher Scientific) as indicated by the manufacturer. Non-transfected receptor cells were stained with CellTrackerTM CMFDA Green (Thermo Fisher Scientific) which is well retained through generations from mother to daughter cells but not to adjacent cells. Twenty-four h post-transfection, the transfected donor cells were co-incubated overnight with the CMFDA-labelled receptor cells and subsequently fixed with 4% PFA. For microscopy analysis, cells were then permeabilized with 0.1% Triton X-100, and stained with DAPI and N protein-specific MAb using with goat anti-mouse IgG conjugated to Alexa Fluor 555 (1:1,000) (Thermo Fisher Scientific). Immunofluorescence images were acquired through an epifluorescence microscope (Olympus BMAX51) equipped with a ProgRes C3 camera (Jenoptics). For quantification, the percentage of N-positive cells for both fluorophores, CMFDA (green) and N protein or mCherry (red) with respect to the total N protein or mCherry plasmid DNA transfected cells was calculated. Inhibition studies of intercellular N protein transfer were performed by incubating cells with Brefeldin A (10 µg/ml), Bafilomycin A1 (100 nM) or VPS-34 inhibitor (1 µM) for 2 h at 37°C before fixation; quantification was performed as described above.

### Statistical analysis

The results are derived from at least two independent experiments and presented as the mean with standard error of the mean (SEM). Impaired Student t-test was calculated using GraphPad Prism software version 6.0 and P values considered significant as follows: * p<0.05, ** p<0.01, *** p<0.001, **** p<0.0001.

## RESULTS

### Cell surface localization of the N protein in hantavirus-infected cells

First, we aimed to investigate the cellular distribution of the hantavirus N protein in infected Vero E6 cells through immunofluorescence studies under permeabilized and non-permeabilized conditions. To label the plasma membrane, we used the membrane probe CellMask^TM^ Orange, an amphipathic membrane dye that is resistant to fixation but dissipates upon treatment with detergents used in cell permeabilization. In permeabilized ANDV-infected cells, N protein-staining was associated with round and filamentous patterns of diverse sizes in the cytoplasm and absence of the plasma membrane dye (Fig. 1A). Strikingly, in non-permeabilized ANDV-infected cells, N protein-staining localized to the plasma membrane, suggesting that a proportion of the N protein was distributed to the cell surface (Fig. 1B). Under the non-permeabilized condition, the N protein appeared mainly round of different sizes rather than filamentous and was dispersed throughout the cell surface (Fig 1B).

**Fig 1.**
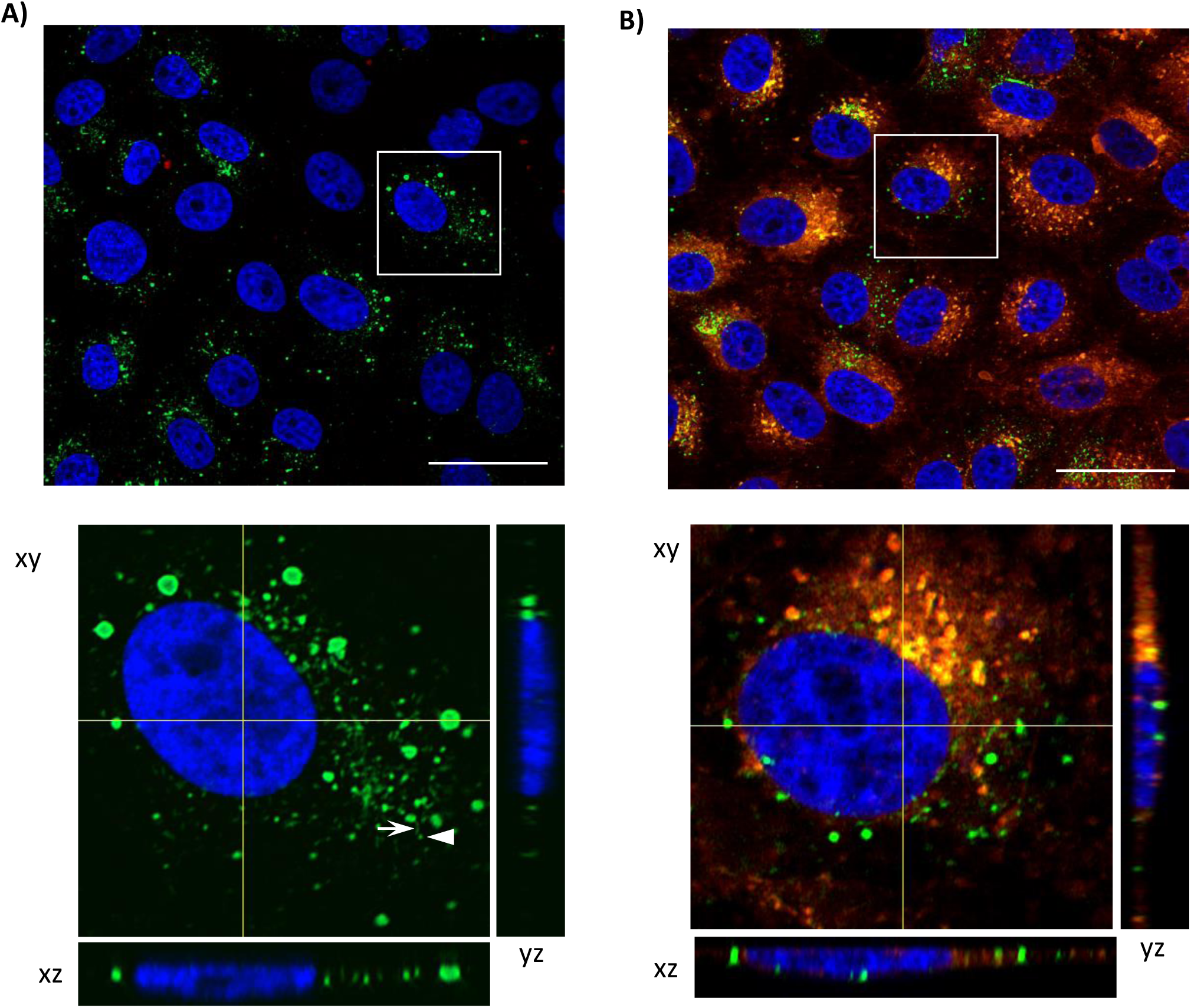
ANDV N surface localization on infected Vero E6 cells. **A-B)** Top panels show representative fluorescence confocal microscopy images of **A)** permeabilized and **B)** non-permeabilized cells infected with ANDV (MOI 1). 24 hpi cells were subjected to Live/Dead staining, fixed with PFA and labelled either directly with anti-N MAb and plasma membrane dye for cell surface staining, or after permeabilization with Triton X-100 for whole cell staining. Bottom panels indicate an optical zoom of the selected cell from the top panel. Orthogonal views were prepared with Fiji software to visualize spatial distribution. The confocal images show the xy plane (main panel), and orthogonal yz and xz projections (side and bottom lateral panels). ANDV N is labelled in green (Alexa Fluor 488), plasma membrane in orange (CellMask^TM^ Orange), nuclei in blue (DAPI). In the bottom panel A, white arrow exemplifies filamentous N-staining, white arrow head shows a round N-staining. Scale bar 100 μM. Images representative of >50 analyzed cells.

Next, we performed flow cytometry of ANDV- or TULV-infected Vero E6 cells. We collected cells at 18, 24, and 48 hpi and stained them with the Live/Dead label to analyze only live cells with intact plasma membrane integrity. Subsequently, we fixed the cells for virus inactivation and next labelled for N protein, with or without prior permeabilization step. Flow cytometry results revealed a time-dependent detection of the ANDV N protein in 38% to 61% for permeabilized ANDV-infected Vero E6 cells (Fig. 2A and 2B). In non-permeabilized ANDV-infected cells, N protein was detected at all examined infection time points, reaching 17% at early time points and 36% at later time points (Fig. 2B). When normalized to total infected cells, the percentage of cells bearing N protein on their surface ranged from 48% to 65% at earlier and later time points post-infection, respectively, without a statistically significant difference (Fig. 2C). As observed for ANDV, the N protein of TULV was readily detected on the cell surface of infected Vero E6 cells over time with concomitant increase of N protein expressing cells and their respective GMFI, ranging from 10% to 25% in non-permeabilized cells (surface) compared to 15% to 38% in permeabilized cells (total) (Fig. 2D and 2E). Referenced to total TULV-infected cells, the percentage of cells carrying N protein on their surface ranged from 57% to 67% at different time points, without statistically significant differences as observed for ANDV-infected Vero E6 cells (Fig. 2F).

**Fig 2.**
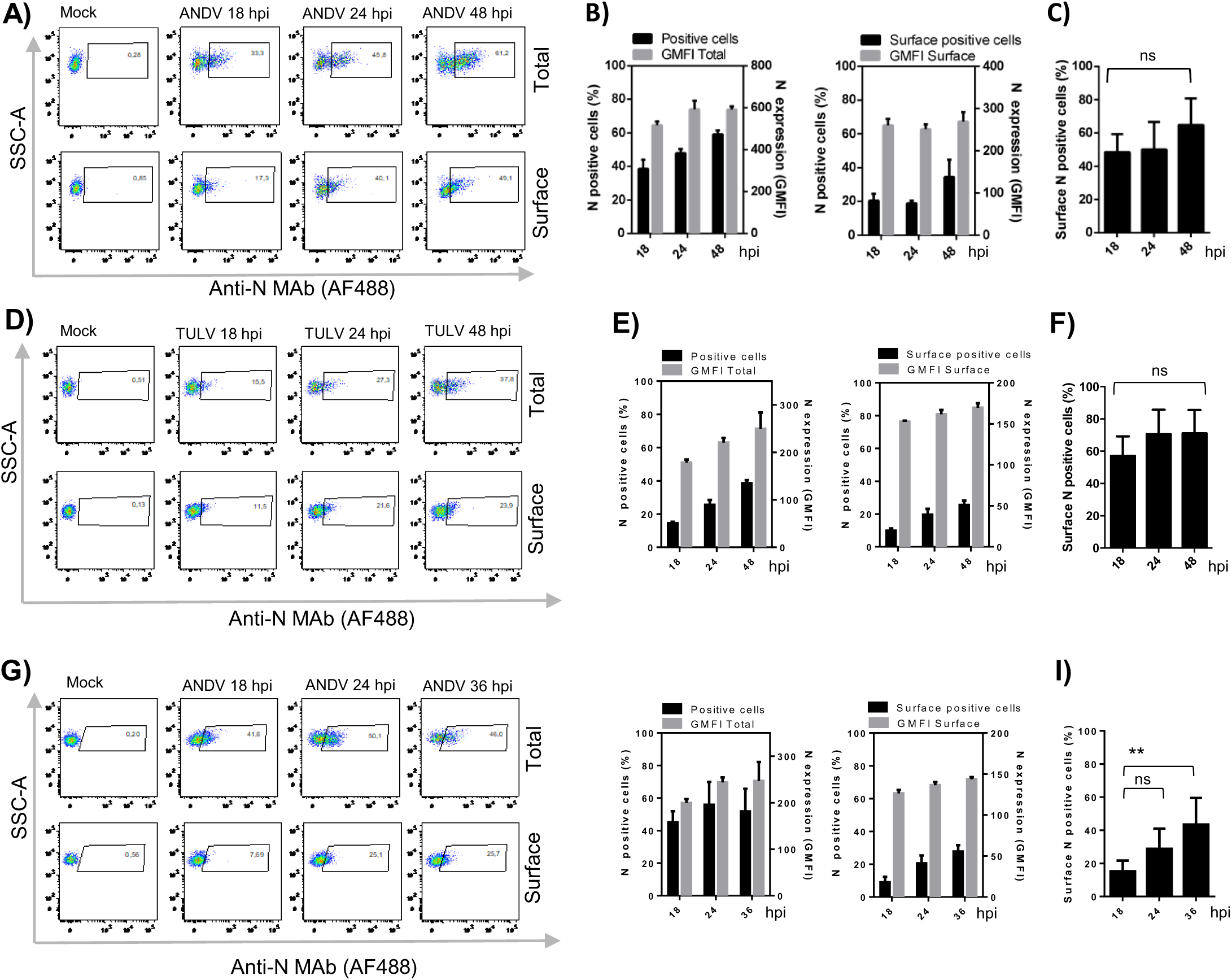
N protein surface localization on cells from different cell lines infected with ANDV and TULV. **A)** Representative dot plots from flow cytometry analysis of N protein labelling of ANDV-infected, permeabilized cells (top panel) and non-permeabilized cells (bottom panel). After 18, 24 and 48 hpi with ANDV (MOI 1), cells were subjected to Live/Dead staining, fixed with PFA and labelled either directly with anti-N MAb (Alexa Fluor 488) for cell surface staining, or after permeabilization with Triton X-100 for whole cell staining. **B)** Quantification of results from (A) of N protein positive, permeabilized (positive cells, left panel) or non-permeabilized ANDV-infected Vero E6 cells (surface positive cells, right panel) indicated in black bars and accompanied with the corresponding geometric mean fluorescence intensity (GMFI) of each condition shown in grey bars. Results in duplicate representative for 2 biological replicates. **C)** Percentage of cells bearing N protein on their surface when normalized to total ANDV-infected cells at 18, 24 and 48 hpi. Results from three biological replicates. **D)** Representative dot plots from flow cytometry analysis of N protein staining in TULV-infected Vero E6 cells (MOI 1) as indicated in (A). **E)** Quantification of results from (D) of N protein positive, permeabilized (positive cells) or non-permeabilized TULV-infected Vero E6 cells as indicated in (B). **F)** Percentage of cells bearing N protein on their surface when normalized to total TULV-infected cells at 18, 24 and 48 hpi as indicated in (C). **G)** Representative dot plots from flow cytometry analysis of N protein staining in ANDV-infected EA.hy926 cells (MOI 1) at 18, 24 and 36 hpi as indicated in (A). **H)** Quantification of results from (G) of N protein positive, permeabilized (positive cells) or non-permeabilized ANDV-infected EA.hy926 cells as indicated in (B). **I)** Percentage of cells bearing N protein on their surface when normalized to total ANDV-infected EA.hy926 cells at 18, 24 and 36 hpi as indicated in (C). The statistical analysis were performed with Student’s t-test; * p < 0.05, ** p < 0.01; *** p < 0.001; ns, not significant.

We also analyzed endothelial cells for N protein surface localization during hantavirus infection by infecting human endothelial EA.hy926 cells with ANDV. Given reduced cell viability due to viral infection, the kinetics assay was reduced to a 36 hpi maximum. As observed in Vero E6 cells, ANDV-infected EA.hy926 cells showed an increase over time in the percentage of N-positive cells, ranging from 9% to 27% and a rise of the respective GMFI of the cells (Fig. 2G and 2H). Different to Vero E6 cells, in EA.hy926 cells we found that the percentage of N protein-expressing cells bearing N protein on their surface increased here from 15% at early to 43% at later time points, revealing a statistical difference between 18 and 36 hpi (Fig. 2I). Collectively, our findings indicate that the N protein reaches the surface of infected cells, independent from the cell type for at least two hantavirus species.

### The localization of the ANDV N protein at the cell surface is independent of other viral components

We next asked whether the localization of the N protein at the cell surface was self-driven, or if it instead relied on other viral components such as the viral Gn/Gc spikes inducing viral assembly and release. To this end, we surveyed HEK293FT cells transfected with a plasmid coding for ANDV N alone, or in combination with a plasmid coding for ANDV GPC. At 48 hpi we labelled live cells with Live/Dead dye to select cells with an intact plasma membrane integrity. Next, we fixed the cells and stained with anti-N or anti-Gc MAb with our without prior permeabilization and acquired the signals by flow cytometry. As expected, we found a Gc signal in permeabilized and non-permeabilized cells, and in presence or absence of N protein co-expression, the GMFI did not vary significantly (Fig. 3A and 3B). The percentage of Gc-expressing cells was further equivalent to the rate of cells bearing Gc on their surface (11% in the presence or absence of N protein) (Fig. 3C), indicating that Gc reaches the cell surface in nearly all cells expressing Gc. In cells expressing either N protein alone or together with GPC, we could also detect N at the cell surface in non-permeabilized cells, (Fig. 3D and 3E), consistent with our results obtained from viral infection (see Fig 2A and 2B). Similar to the detection of Gc, surface N protein was present in most N-expressing cells, suggesting that in the absence of the viral envelope proteins, N protein arrives at the cell surface of virtually all HEK293FT cells in which it is being produced (Fig. 3F). We repeated the transfection experiment also in Vero E6 and EA.hy926 cells. Despite a lower transfection efficiency in these cells reflected in the lower percentage of permeabilized N-positive cells, the N protein was present on the plasma membrane in 46% of N-expressing Vero E6 cells and 62 % of N-expressing EA.hy926 cells with no significant differences when GPC was co-expressed (Fig. S1).

**Fig 3.**
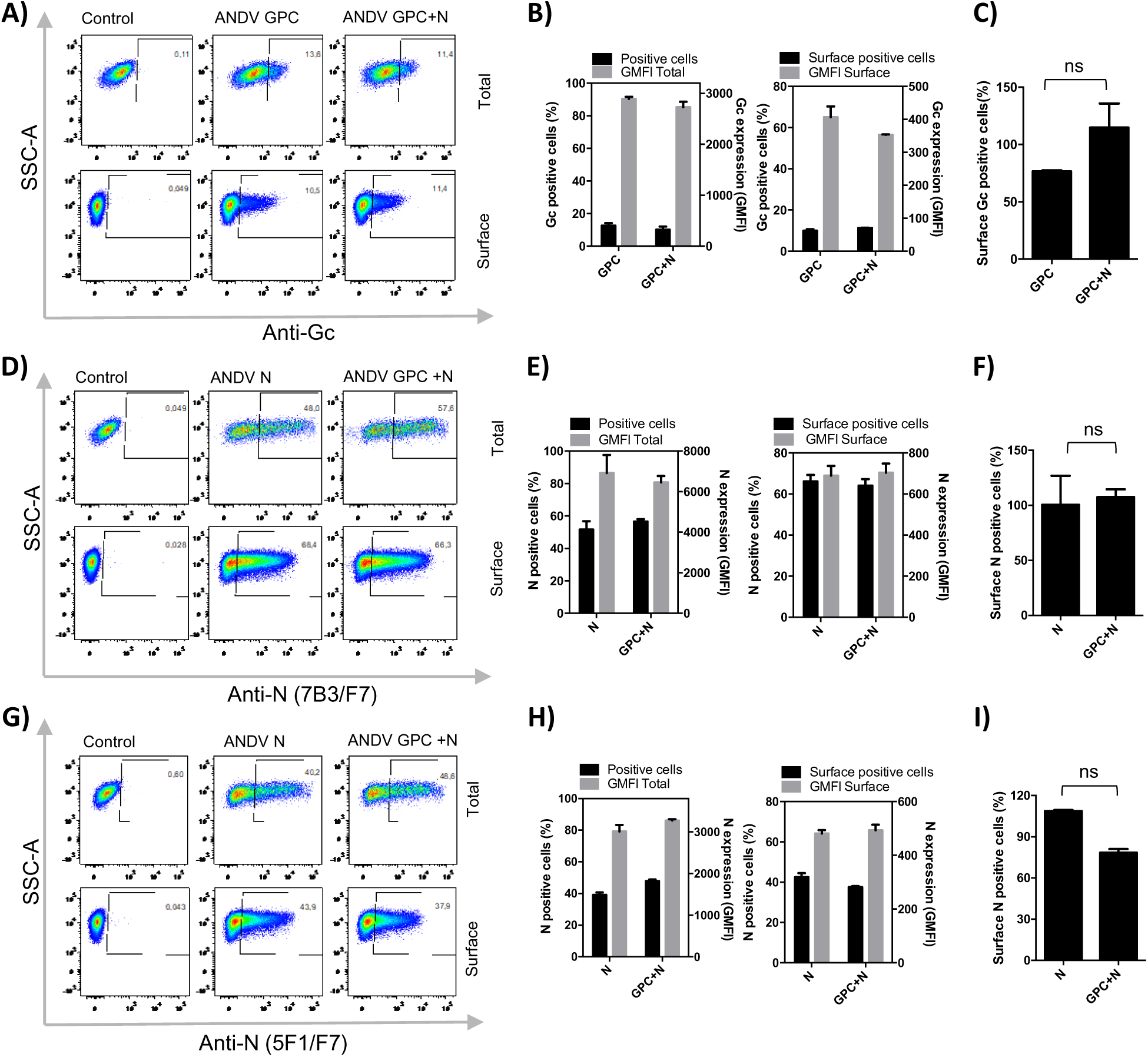
The ANDV N protein cell surface localization is independent from Gn/Gc spike-mediated viral assembly. **A)** Representative dot plots from flow cytometry analysis of Gc protein labelling of transfected, permeabilized (top panel) and non-permeabilized HEK293FT cells (bottom panel). HEK293FT cells were either transfected with ANDV GPC plasmid encoding Gn/Gc, co-transfected with ANDV GPC plasmid and ANDV N-encoding plasmid or an empty control plasmid (pI.18). 48 h post-transfection, cells were subjected to Live/Dead staining, fixed with PFA and labelled with anti-Gc MAb (Alexa Fluor 488) for cell surface staining, or after permeabilization with Triton X-100 for whole cell staining (total). **B)** Quantification of results from (A) of Gc protein positive, permeabilized (positive cells, left panel) or non-permeabilized HEK293FT cells (surface positive cells, right panel) expressing Gn/Gc or Gn/Gc and N protein. The percentage of Gc-positive cells is indicated in black bars and each condition is accompanied with the corresponding geometric mean fluorescence intensity (GMFI) shown in grey bars. **C)** Percentage of HEK293FT cells bearing Gc protein on their surface when normalized to total Gc-expressing cells. **D)** Representative dot plots from flow cytometry analysis of N protein labelling in HEK293FT cells transfected with ANDV N coding plasmid or ANDV N and ANDV GPC coding plasmids. **E)** Quantification of results from (D) of N protein positive, permeabilized (positive cells) or non-permeabilized transfected HEK293FT cells (surface positive cells) expressing N protein or N and Gn/Gc proteins. The percentage of N-positive cells is indicated in black bars and GMFI in grey bars. **F)** Percentage of cells bearing N protein on their surface when normalized to total N protein expressing cells. **G)** Representative dot plots from flow cytometry analysis of N protein labelling with alternative MAb (clone 5F1/F7) in HEK293FT cells transfected with ANDV N coding plasmid or ANDV N and ANDV GPC coding plasmids. **H)** Quantification of results from (G) as indicated in (E). **I)** Percentage of cells bearing N protein on their surface when normalized to total N protein expressing HEK293FT cells using MAb clone 5F1/F7 for N protein labelling. **(B, E, H)** Results in duplicate representative for 2 biological replicates. **(C, F, I)** Results from three biological replicates. Statistical analysis were performed with Student’s t-test; * p < 0.05; ** p < 0.01; *** p < 0.001; **** ns, not significant.

To rule out the possibility that the detected N protein on the cell surface of cells expressing N protein alone was an artefact produced by the used Mab (clone 7B3/F7), we performed the same assay with an alternative anti-N MAb (clone 5F1/F7), known to target to a different epitope on N protein [37]. As before, using this antibody we detected a similar cell surface localization of N protein in the presence or absence of GPC co-expression; with an average of 65% and 79%, respectively (Fig. 3G-3I). Together, the results indicate that expression of ANDV N protein leads to its arrival on the extracellular face of the plasma membrane by an unknown mechanism that is independent of the viral Gn/Gc glycoproteins or other viral components.

### The N protein of various hantaviruses localizes to the cell surface and is secreted into the supernatant

Next, we wondered if cell surface localization was a unique feature of the ANDV and TULV N proteins. Therefore, we transfected Vero E6 cells with plasmids coding for the N proteins of ANDV, SNV, TULV and PUUV. Using the same experimental approach, the transfection efficiency of plasmids coding for N protein from the different hantavirus species varied strongly, yielding N-expression in permeabilized cells ranging from 19% to 58% (Fig. 4A and 4B). Interestingly, we detected the N protein of all tested hantaviruses on the plasma membrane, with ANDV N showing the highest and PUUV N protein the lowest ratio of surface localization ranging from 25% to 62% of N-expressing cells (Fig. 4C).

**Fig 4.**
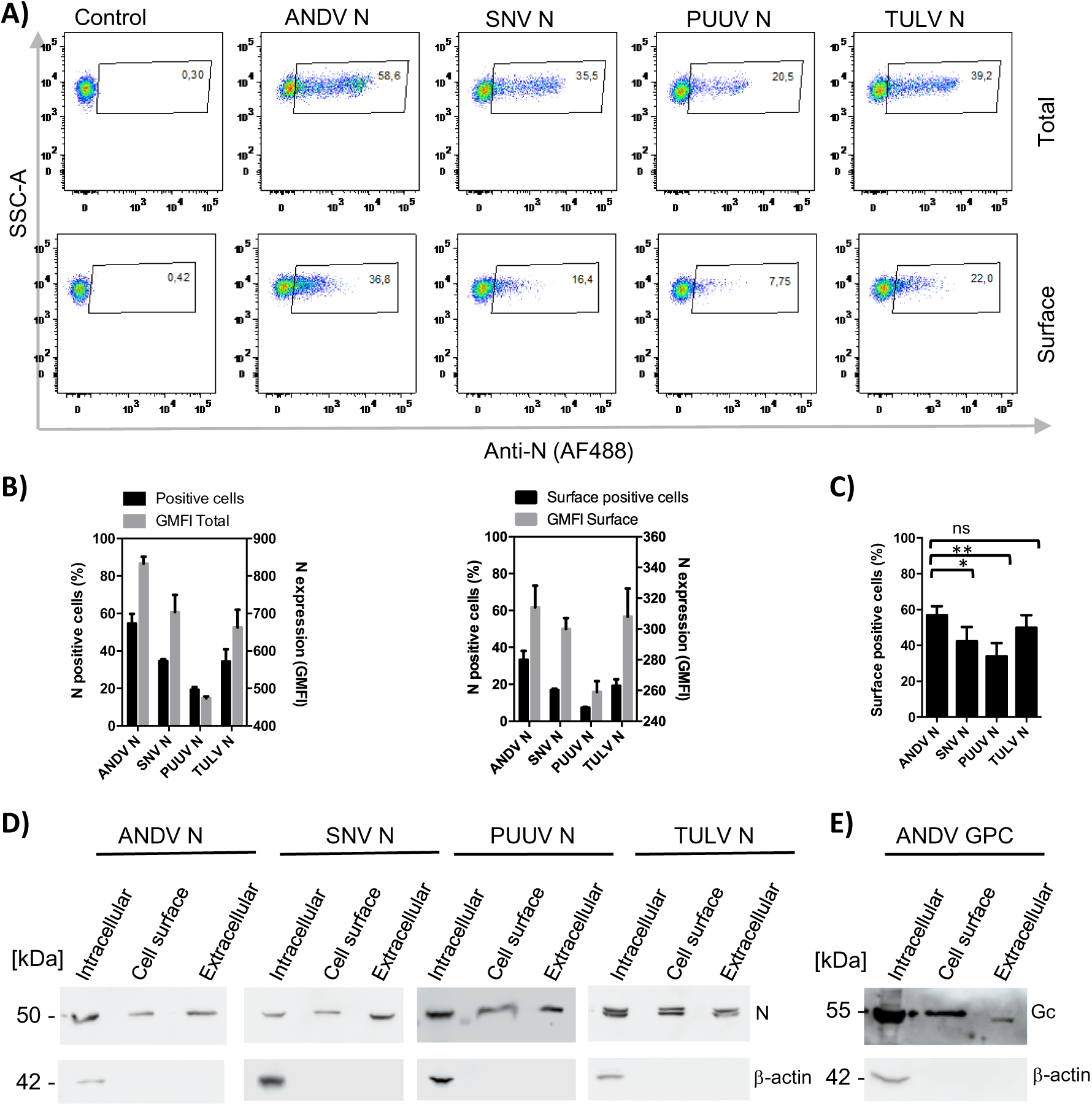
Surface localization and extracellular secretion of N protein from different hantaviruses. **A)** Representative dot plots from flow cytometry analysis of N protein labelling of transfected, permeabilized (top panel) and non-permeabilized Vero E6 cells (bottom panel) transfected with plasmids encoding N protein from ANDV, SNV, PUUV or TULV, or empty control plasmid. 48 h post-transfection, cells were subjected to Live/Dead staining, fixed with PFA and labelled with anti-N MAb (Alexa Fluor 488) for cell surface staining, or after permeabilization with Triton X-100 for whole cell staining (total). **B)** Quantification of results from (A) of N protein positive, permeabilized (positive cells; left panel) or non-permeabilized transfected Vero E6 cells (surface positive cells, right panel) expressing N protein. The percentage of N-positive cells is indicated in black bars and geometric mean fluorescence (GMFI) in grey bars. **C)** Percentage of cells bearing N protein from different hantaviruses on their surface when normalized to total N protein expressing cells from the corresponding virus. **D)** Western blot analysis of intracellular fraction, cell surface fraction and cell supernatant samples of HEK293FT cells expressing N protein from different hantaviruses. Hek293FT cells transfected with different hantavirus N expression plasmids and 48 h later the cell supernatant was collected and total proteins precipitated using ammonium sulfate. The cells were biotinylated at 4°C and after lysis, the cell surface fraction (biotinylated) separated from intracellular proteins. N proteins were detected by anti-N MAb (clone 7B3/F7) and anti-β-actin MAb used as control of cellular integrity. **E)** Western blot analysis of intracellular fraction, cell surface fraction and cell supernatant samples of HEK293FT cells expressing Gn/Gc proteins from ANDV. Hek293FT cells were transfected with ANDV GPC plasmid coding for Gn/Gc spikes, the samples prepared as in (D) and Gc detected with anti-Gc MAb (clone 2H4/F6).

To further confirm our observations by fluorescence microscopy and flow cytometry of non-permeabilized cells, we separated intracellular and surface proteins experimentally by biotin cell surface labelling. For this, cell surface proteins were specifically biotinylated and independently recovered through neutravidin agarose purification. In this experimental setup, ANDV GPC was used as a positive control. In cells transfected with the different N protein-encoding plasmids, the western blot analysis using a specific anti-N MAb revealed a clear migration species of approximately 50 kDa in intracellular fractions, corresponding to the expected molecular weight of N protein (Fig. 4D). Only in the case of TULV N protein, a double band was observed in all experimental conditions; which may be caused by post-translational modifications. Confirming our previous observations, the viral N protein was also detected in the biotinylated cell surface fractions (Fig. 4D). As expected, the positive control Gc, which has been extensively characterized for its presence on the plasma membrane was also detected in both analyzed fractions (Fig. 4E) [63, 64]. Additionally, β-actin, which is located juxtaposed below the plasma membrane, was labelled as a control. A β-actin signal was only evident in the intracellular fractions, and not in fractions corresponding to the cell surface. This differential signal confirms the integrity of the cells during the biotinylating procedure and affirms the specificity of the experiment. Together, the multiple experimental approaches, including fluorescence microscopy, flow cytometry and specific cell surface protein labelling, confirm the presence of the N protein from all tested hantaviruses on the cell surface of infected or transfected cells, suggesting a common trait (Fig. 4A-4D).

The observation of N protein localization on the plasma membrane prompted us to investigate whether the hantavirus N protein is also secreted into the extracellular environment. To test this, we used ammonium sulfate to precipitate the total protein content of the cell supernatants from transfected cells and tested for the presence of N protein by western blot analysis. By using this approach, we readily detected N protein in the precipitates of all supernatants from cells expressing the different hantavirus N proteins (Fig. 4D), as observed for the viral Gc protein (Fig. 4E) which is known to be secreted from cells in the form of VLPs [46]. As expected, the negative β-actin control was not detected in the supernatant of the cells, confirming that the presence of hantavirus N proteins in the cell culture supernatant was not attributable to cellular damage.

### ANDV N protein is transferred to neighboring cells

Several viral proteins are known to be secreted into the extracellular space and further transferred to neighboring cells, such as SARS-CoV-2 N protein, dengue virus NS1 and HIV-1 Tat [50, 65, 66]. Based on these observations, we analyzed the possible transfer of the N protein from N-expressing donor cells to non-expressing neighboring cells. To distinguish receptor cells from N-expressing donor cells, receptor cells were labelled with CMFDA before their co-culture. Donor cells were either transfected with the ANDV N expression plasmid or with a control plasmid coding for the cytoplasmic mCherry protein. Twenty-four h post-transfection the transfected donor cells were co-cultured with CMFDA-labelled receptor cells for 24 h. For immunofluorescence analyses, the cells were subsequently fixed, permeabilized and incubated with anti-N protein MAb (Fig. 5A). In CMFDA-labelled cells co-cultured with mCherry-expressing cells, an average of 2.6% cells stained positive for both fluorophores (mCherry and CMFDA), reflecting the low leakiness of the system (Fig 5B). By contrast, when co-cultured with cells expressing the ANDV N protein, an average of 15% of cells were ANDV-N/CMFDA double positive, revealing a significant difference with the mCherry control (Fig. 5B) Interestingly, several double positive (recipient) cells showed a predominant peri-nuclear staining for N, similar to what is observed in N-transfected or ANDV-infected cells (Fig. 5B). A similar experiment was conducted to evaluate the transfer of TULV N protein to neighboring cells. Unlike the ANDV N protein, no significant transfer of the TULV N protein (average of 3%) compared to mCherry control plasmid could be documented (Fig. 5B).

**Fig 5.**
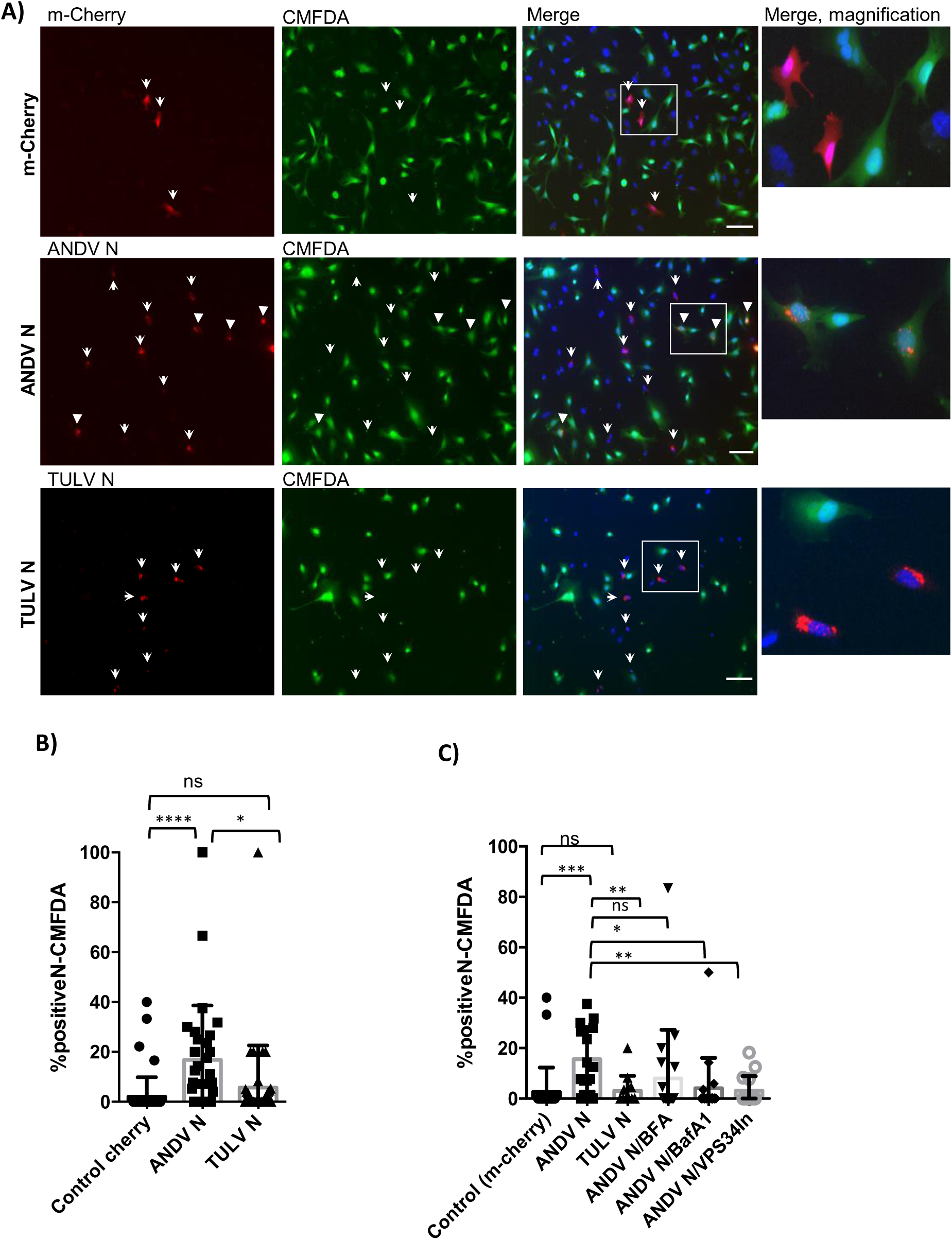
N protein transfer to neighboring cells. A) Representative immunofluorescence microscopy images of co-cultured EA.hy926 cells transiently expressing m-Cherry as control (top panels), ANDV N protein (middle panels) or TULV N protein (lower panels), with CMFDA-non transfected EA.hy926 cells. 24 h after transfection of cells they were detached and co-incubated over-night with CMFDA-labelled EA.hy926 receptor cells. Once fixed with PFA, N-expressing cells were stained with anti-N protein MAb (Alexa Fluor 555). On the right side, magnified image from square from fluorescence merge images are shown. Scale bar, 100 µM. White arrows show N protein or mCherry-expressing cells, while solid arrow heads show N-transfer events of N- and CMFDA-double positive cells. **B)** Quantification of protein transfer to CMFDA-positive receptor cells. **C)** Quantification of protein transfer after inhibitor treatment of cells. Inhibitors were incubated 2 h before fixation, including Brefeldin A (BFA), Bafilomycin A1 (BafA1) and VPS-34 inhibitor (VPS34In)). For quantification, at least 200 cells were analyzed for each condition (from two biological replicates). Statistical analysis was performed with Student’s t-test; * <0.05 ,** P < 0.01; *** P < 0.001; **** P < 0.0001; ns, not significant.

### Inhibition of the ANDV N protein transfer to neighboring cells

To gain further information regarding the pathway by which ANDV exits the cells, the effect of inhibitors of the classical and unconventional secretion pathway on N-transfer to neighboring cells was evaluated. Brefeldin A, the autophagosome formation inhibitor Vps34, and lysosomal acidification inhibitor Bafilomycin A1 were used as secretion inhibitors [67]. For this assay, transfected cells were co-cultured as described above and subjected to inhibitor treatment 2 h before fixation. Quantification of ≥200 cells per condition, demonstrated a significantly reduced ANDV N protein signal in the CMFDA-labeled receptor cells, for all the inhibitors analyzed with less significance for Vps34 (Fig. 5C). Thus, our results provide evidence that the ANDV protein is transferred to neighboring cells, which can be reduced by blocking specific secretory pathways; yet the precise secretion pathway(s) and cell entry mechanisms used by the ANDV N protein remain to be unveiled.

## DISCUSSION

To complete their replication cycle, viruses have evolved diverse strategies to exploit the host cellular machinery and suppress antiviral responses. Because viral genome size limits the number of encoded proteins, viruses with small genomes often rely on multifunctional proteins to perform a range of essential roles. The hantavirus N protein exemplifies this versatility: in addition to mediating genome encapsidation, it is crucial for viral RNA transcription and translation, as well as for inhibiting apoptosis and antagonizing the host antiviral innate immune responses in the cytoplasm [14–18, 26, 68–70].

Here, we demonstrate that, contrary to previous descriptions of the hantavirus N protein as being confined to the cytoplasm or incorporated into enveloped virions, N proteins from multiple hantavirus species are also present on the surface of various infected cell types, including endothelial cells. Moreover, our data indicate that the N protein is secreted into the extracellular medium and transferred to neighboring, uninfected cells. This finding aligns with the strong and persistent anti-N humoral immune response against the N protein observed in infected patients and rodent hosts [32, 33, 36–38, 44]. Despite reports of N protein secretion in several other (-)ssRNA viruses [71], the literature has not previously recognized the hantavirus N protein, or any N protein from members of the *Bunyaviricetes* class [72] as secretory proteins.

The data presented here show that the N protein of hantaviruses from various geographic regions and four different rodent host subfamilies does not require the presence of any other viral component to reach the plasma membrane and translocate across a cellular membrane, indicating that intrinsic properties of the protein mediate these processes. The molecular mechanisms driving N protein localization to the cell membrane and exit remain to be elucidated. However, it is tempting to speculate that this process likely occurs via an unconventional secretory pathway [73], since the N protein of hantaviruses, and of RNA viruses in general, lack a classical hydrophobic signal sequence that would enable their recognition by the signal recognition particle and co-translational translocation into the lumen of the endoplasmic reticulum [74]. Therefore, an unconventional protein secretion (UPS) pathway may be involved [73], most likely through direct protein translocation across a cellular membrane (type I or II), as described for the human immunodeficiency virus Tat protein [66, 75], or through a vesicle-mediated export mechanism (type III or IV).

Our results suggest that the N protein reaches the extracellular space in a non-enveloped, free form, as indicated by its direct detection with different anti-N MAbs. Such binding would have been impeded if the N protein were enclosed within lipid membranes, for example, inside exosomes or viral particles. The surface localization and secretion of N protein in the absence of other viral components, such as viral RNA, further indicate that at least a fraction of the heterologously expressed N protein is not incorporated into viral ribonucleocapsids as characterized previously [20]. Although BFA treatment reduces cell-surface localization of the N proteins of measles virus and RSV [51, 52], such an effect may reflect alterations in intracellular trafficking of the N protein rather than a direct inhibition of its release. When we treated infected cells with drugs that block conventional or unconventional secretory pathways, all compounds reduced N protein transfer to neighboring cells. Given this reduction, we propose that these inhibitors may indirectly interfere with the transport of host cellular factors required for N protein secretion and intercellular transfer. Additionally, we cannot exclude the possibility that N protein transfer occurs through direct cell–cell contacts.

The cell-surface localization and secretion of N proteins from RSV, measles virus, and human coronaviruses (CoVs) have been associated with key functions during infection, particularly in promoting immunosuppression through multiple mechanisms. The RSV N protein impairs antigen presentation by disrupting immunological synapse formation, thereby inhibiting T cell activation [52]. The measles virus N protein binds FcγRII on B cells, reducing immunoglobulin synthesis and suppressing interleukin (IL)-12 secretion, cell proliferation, and cytokine responses [51, 76, 77]. The N protein of SARS-CoV-2 and other human CoVs binds heparan sulfate and various chemokines, disrupting leukocyte chemotaxis [50, 78] and facilitating intercellular N protein transfer, collectively contributing to immune evasion and modulation. Distinct immunomodulatory roles have also been reported for the hantaviral N protein, including the inhibition of apoptosis and interference with intracellular signaling pathways, such as the Jak/STAT and NF-κB pathways, as well as direct caspase inhibition [26, 28, 79]. Thus, the newly incorporated N protein could prime uninfected cells, targeting them for infection, as in the presence of the N protein, cells would be limited in their ability to mount an efficient antiviral response [26, 28, 79]. In addition, the N protein may also modulate apoptosis on the cell surface of infected cells in the extracellular environment, for instance by interfering with granzyme B activity during cytotoxic lymphocyte–mediated cell lysis [28]. In this line, one study suggests that PUUV–infected endothelial cells are protected from natural killer (NK) cell-mediated cytolysis, whereas uninfected neighboring cells are not [80]. The findings presented here, showing that the hantavirus N protein localizes on the cell surface and is transferred to neighboring cells, raise the question of whether NK cells could be activated through a non-spike antibody-specific mechanism. Such a pathway could also explain the strong activation of CD56dim NK cells observed in HFRS patients [80]. In the case of SARS-CoV-2, non-spike antibody-dependent NK cell activation has been shown to play a crucial role in antiviral immunity and vaccine-induced protection across multiple viral strains [81]. In COVID-19 patients, the detection of free N protein in serum and urine has been correlated with disease severity [82, 83]. In our study, we observed significant differences in the transfer efficiency of the N protein from ANDV compared with that of the non-pathogenic TULV. The enhanced ability of ANDV N protein to transfer between cells may represent a contributing factor to its pathogenic potential. However, further investigation is required to elucidate the underlying mechanisms and implications. Moreover, previous studies have shown that the ANDV N protein increases the permeability of microvascular endothelial cells, a characteristic feature of ANDV infection *in vitro*, which may contribute to the vascular leakage observed in patients [29, 84, 85].

Finally, it is important to note that the N protein from various hantaviruses has been evaluated as a vaccine antigen in multiple animal models, where it reduces lethality. This protective effect has been primarily attributed to cellular immune responses, particularly CD8⁺ T cell–mediated immunity and antibody-dependent cellular cytotoxicity [36, 86–89]. These findings are consistent with vaccine studies on SARS-CoV-2 and influenza A virus that include the N protein in combination with spike antigens. In those models, the high immunogenicity and antigenic stability of N protein have made it a strong candidate for broad-spectrum protection, particularly in cross-variant or cross-strain immunity [90–92].

In summary, the data presented in this study reveal novel insights into our fundamental understanding of hantavirus infections and virus-host cell interactions. Although the functional significance of N protein surface localization, secretion and intercellular transfer remains to be determined, it suggests that it has a role in viral dissemination and pathogenesis. The identification of extracellular N protein of hantaviruses, decades after its discovery for other RNA viruses, underscores the need to investigate its functions in the extracellular milieu, and opens new avenues for diagnostic, preventive, and therapeutic strategies. These new perspectives may extend to additional bunyaviruses that pose thread to public health.

## Funding

This study was financed by ANID, through grants FONDECYT 1221811, FONDECYT 11230653, FONDEF TA24I10051 and Basal FB210008.

## Acknowledgements

We thank for the possibility to use the BSL3 facility at the Centro de Investigaciones Médicas (CIM), Facultad de Medicina, Pontifícia Universidad Católica de Chile. We also thank Dr. Héctor Galeno from Instituto de Salud Pública de Chile for providing the ANDV strain CHI-7913 and Drs. Detlev H. Krüger and Boris Klempa (Charité Universitätsmedizin Berlin, Germany) for providing TULV, strain Moravia/5302Ma/94. We are furthermore thankful to Dr. Jim Robertson (National Institute for Biological Standards and Control, Hertfordshire, UK) for providing the pI.18 vector. The technical assistance of Anna Hegele and Beate Becker Ziaja in the cloning of the PUUV and TULV sequences is kindly acknowledged.

---

**Fig S1.**
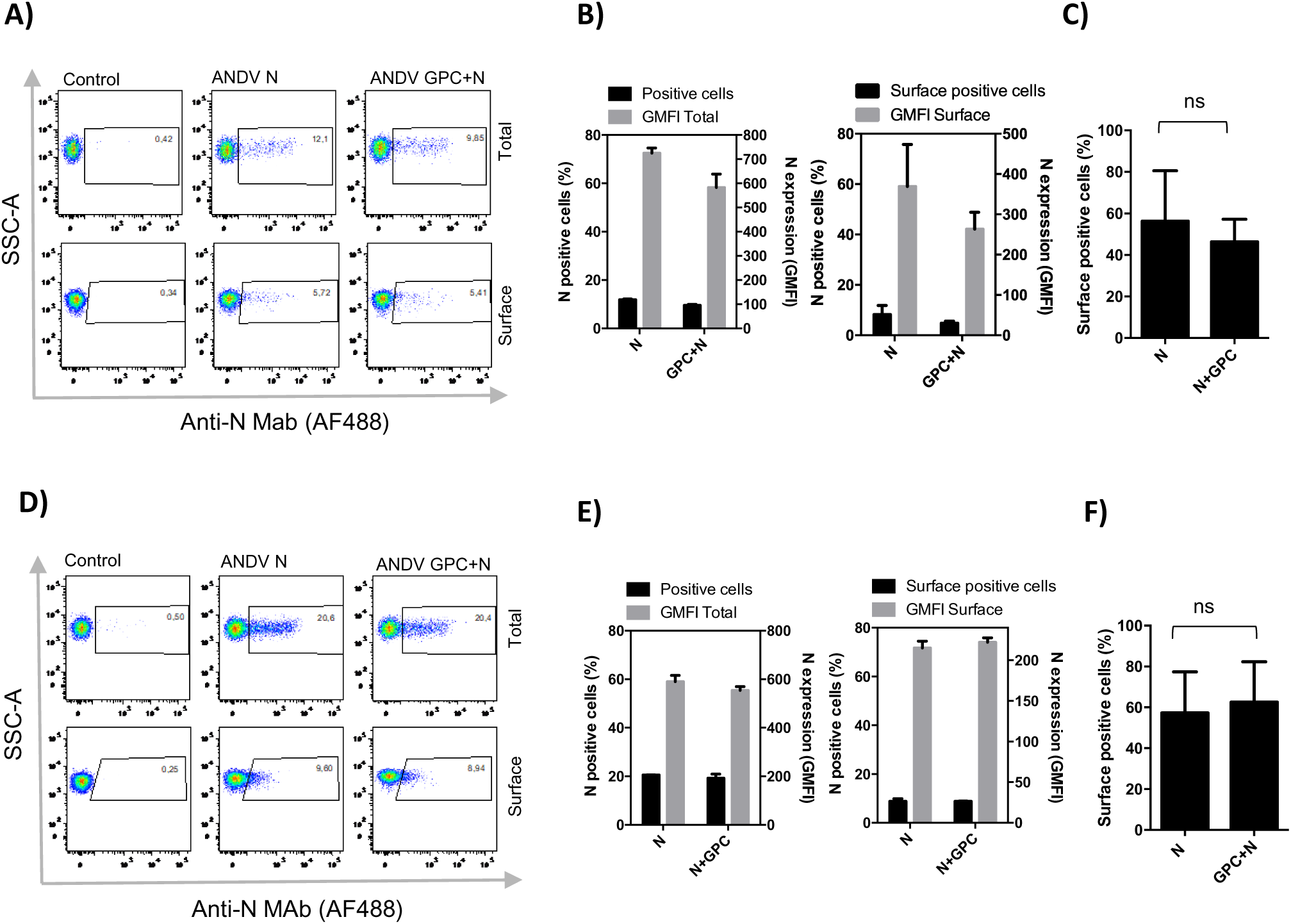
Different cell lines transfected with ANDV N protein show N protein cell surface localization independent from Gn/Gc spike-mediated viral assembly. **A, D)** Representative dot plots from flow cytometry analysis of total (top panel) and surface (bottom panel) of Vero cells **(A)** or EA.hy926 cells **(D)** transfected with control plasmid, ANDV N coding plasmid or ANDV N and ANDV GPC coding plasmids. 48 h post-transfection, cells were stained with Live/Dead dye and incubated with anti-N Mab (clone 7B3/F7) with or without permeabilization. **B, E)** Quantification of results from (A or D) of N protein positive, permeabilized (positive cells, left panel) or non-permeabilized transfected HEK293FT cells (surface positive cells, right panel) expressing N protein or N and Gn/Gc proteins. The percentage of N-positive cells is indicated in black bars and GMFI in grey bars. Results in duplicate representative for 2 biological replicates. **C, F)** Percentage of cells bearing N protein on their surface when normalized to total N protein expressing cells. Results from three biological replicas. Statistical analysis were performed with Student’s t-test; * p < 0.05, ** p < 0.01; *** p < 0.001; **** p < 0.0001; ns, not significant.

## Notes

### Competing Interest Statement

The authors have declared no competing interest.

